# A Human Vocal Fold Organ-On-Chip for Studying Platform-Dependent Mucosal Responses to Particulate Matter

**DOI:** 10.64898/2026.05.29.728871

**Authors:** Patrick T. Coburn, Meghana Munipalle, Yin Liu, Vlasta Lungova, Samjhana Thapa, Lisa Martignetti, Xinyu Liu, Gilles Maussion, Carol X.-Q. Chen, Thomas M. Durcan, Susan L. Thibeault, Nicole Y. K. Li-Jessen

## Abstract

**Background:** Coarse particulate matter (PM_10_) deposits at the vocal fold (VF) mucosa, yet upper airway responses remain poorly characterized. Existing *in vitro* VF models use monocultures that lack the stratified epithelium, lamina propria, and physiological perfusion. We developed a chip-based co-culture model of human VF mucosa and applied it to acute PM_10_ exposure.

**Methods:** Vocal fold organ-on-chip (VF-OOC) paired primary VF fibroblasts with either immortalized laryngeal epithelial cells (iLEC) or induced pluripotent stem cells (iPSC)-derived VF epithelial cells. Transwell and 2D chip cultures were controls. Epithelia matured at an air-liquid interface on fibroblast-embedded collagen over a perfused microchannel. PM_10_ urban dust (0 to 400 µg/mL) was applied for 24 hours. Responses were assessed by histology, immunofluorescence, transmission electron microscopy, qPCR, and ELISA.

**Results:** VF-OOC produced a thicker stratified epithelium with upregulated barrier, mucin, and extracellular matrix genes versus transwell controls. Intercellular junctions and basement membrane matched adult human VF mucosa. PM_10_ remained at the epithelial surface across all doses. Transwell_iLEC_ downregulated basal markers (*TP63, KRT5, KRT14*). VF-OOC_iLEC_ upregulated *MUC1* and *HAS3*, consistent with an adaptive mucosal response. VF-OOC_iPSC_ additionally induced suprabasal, junctional, extracellular matrix, and cytokine genes largely absent in iLEC and transwell formats.

**Conclusions:** VF-OOC reproduced key features of native human VF mucosa and captured distinct, platform-specific responses to PM_10_ that differed by epithelial cell source. By introducing new approach methodologies (NAMs) into laryngology, this platform extends respiratory toxicology to the upper airway beyond bronchial and alveolar compartments and allows mechanistic studies of exposure-linked diseases such as laryngitis.

## 1. INTRODUCTION

Particulate matter (PM) air pollution is associated with 4.7 million premature deaths per year and contributes to 8% of global disability-adjusted life years [1, 2]. PM is a heterogeneous mixture of heavy metal ions, dust, polycyclic aromatic hydrocarbons, pollen, and microbial agents from vehicular emissions, industrial operations, and wildfires [3, 4]. While PM exposure has been extensively studied in the context of asthma, chronic obstructive pulmonary disease, and lung cancer [1, 5], its effects on the upper airway remain poorly characterized. Other inhaled exposures with a particulate composition similar to ambient PM, such as wildfire smoke, occupational dust, and e-cigarette aerosols, also affect upper airway health.

Upon inhalation, PM generates excessive reactive oxygen species that activate NF-κB and MAPK signaling and induce pro-inflammatory cytokines such as interleukin (IL)-1β, IL-6 and IL-8 [6-8]. PM also degrades tight junction proteins, which increases epithelial permeability and compromises barrier integrity [7, 9]. These mechanisms are well characterized in bronchial and alveolar epithelia. Whether intact VF mucosa mounts similar or distinct responses or whether the stratified epithelial barrier attenuates these responses is unknown.

The larynx is the narrowest segment of the respiratory tract and a primary site of coarse PM (PM_10_) deposition [10, 11]. High airflow velocity and anatomical restriction create turbulence that promotes particle deposition on vocal fold (VF) surfaces [12, 13]. Human VF possesses a complex multi-layered, multi-cellular mucosal architecture and immunity [14-20]. The VF mucosa consists of a non-keratinized stratified squamous epithelium overlying a basement membrane and lamina propria populated by fibroblasts in extracellular matrix (ECM) [21, 22]. Tight junctions and adherens junctions seal adjacent epithelial cells and protect the underlying connective tissue from environmental insults [21]. Epidemiological data have linked environmental irritant exposure to chronic laryngitis [23], vocal cord dysfunction [24], and laryngeal cancer [25]. PM_10_ exposure is also associated with increased emergency admissions for laryngitis at a lag of five days [26].

The molecular bases of PM_10_-related VF mucosal injury are poorly understood because existing preclinical models are insufficient for environmental toxicology studies. Previous *in vitro* studies exposed VF fibroblasts in two-dimensional (2D) monocultures to PM for acute toxicity assessment [27, 28]. At concentrations as low as 20 µg/mL, PM induced NLRP3-mediated pyroptosis in VF fibroblast monocultures [29]. The predictive value of monoculture systems is, however, limited because they bypass the stratified epithelial barrier and the cell-cell interactions that govern *in vivo* responses to inhaled particles [21, 30, 31]. An air-liquid interface (ALI) model of primary mouse laryngeal epithelial cells incorporated this epithelial barrier and reproduced cigarette-smoke-induced disruption and recovery [32]. Even so, this ALI model relied on an epithelial monoculture in static transwell inserts and has not been applied to particulate matter or extended to a co-cultured human VF mucosa.

Cell source is another limitation. The only commercially available human laryngeal epithelial cell line was isolated from the posterior glottis and has limited capacity for terminal differentiation [33]. Induced pluripotent stem cells (iPSC)-derived VF epithelial cells are a viable option that recapitulates the full differentiation program of native VF mucosa, including suprabasal markers absent in immortalized lines. These iPSC-derived cells have been combined with VF fibroblasts in a 3D co-culture model to study smoke and e-cigarette vapor responses [34, 35]. Yet, VF epithelial cells have not been tested in organ-on-chip (OOC) platforms, the microengineered cell culture devices that recapitulate native tissue microenvironments through continuous perfusion, physiologically relevant cell-to-media ratios, and shear forces relevant to cell differentiation and gene expression [36-38].

Being one of the new approach methodologies (NAMs), OOC are human-relevant *in vitro* models that the U.S. FDA and European EMA prioritized for reducing animal testing in biomedical research [39-42]. Organ chips have already made substantial advances in modeling the airway [43-50]. For instance, OOC systems have recapitulated cytokine hypersecretion of COPD and asthma exacerbations [51, 52] and replicated pathogen-induced inflammatory responses [53]. Conversely, OOC technology has not been applied to the upper airway or VF tissue.

In this study, we developed and characterized a vocal fold organ-on-chip (VF-OOC) through stepwise optimization of culture dimensionality, physiological scaling, perfusion and fluidic shear (**Figure 1**). Our VF-OOC integrated human primary VF fibroblasts and epithelial cells in a 3D co-culture with ALI and continuous perfusion. To test the utility of the platform for upper airway environmental toxicology, we challenged the microengineered VF mucosae with PM_10_ and compared structural, transcriptional, and inflammatory responses between VF-OOC and conventional transwell cultures. The primary model used immortalized laryngeal epithelial cells (iLEC) and was tested across a PM_10_ dose range for 24 hours. A secondary model using iPSC-derived VF epithelial cells was tested at a selected PM_10_ dose to determine whether cell source influenced the mucosal response to pollutant exposure.

**Figure 1.**
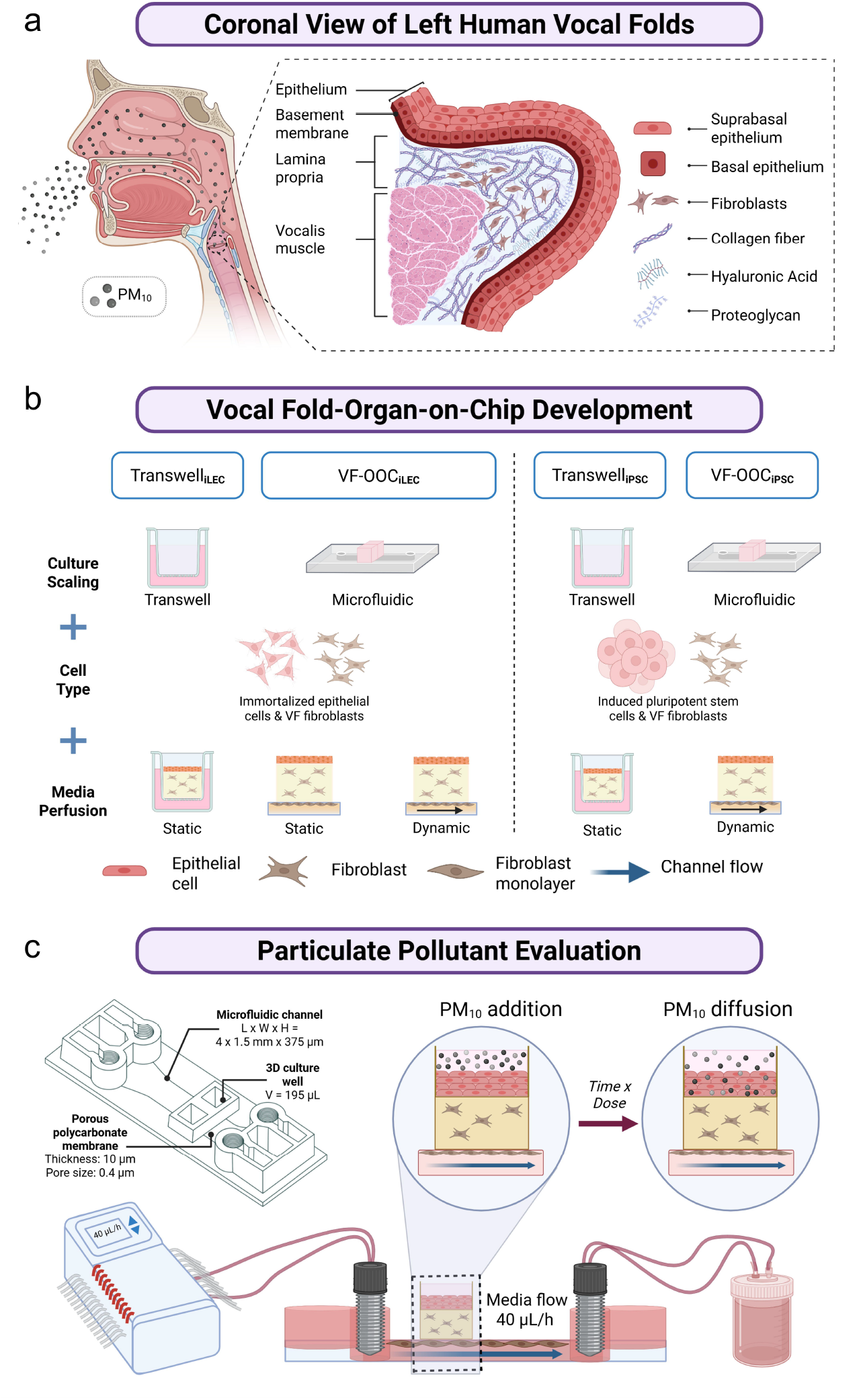
Overview of vocal fold organ-on-chip (VF-OOC) development and PM_10_ evaluation. (a) During breathing, coarse particulate matter (PM_10_) deposits on the vocal fold mucosa. (b) VF-OOC models were developed by systematically varying culture scaling (transwell versus microfluidic chip), epithelial cell source (immortalized laryngeal epithelial cells or iPSC-derived VF progenitors), and media perfusion (static versus dynamic). (c) VF-OOC integrated a 3D epithelial-fibroblast co-culture at an air-liquid interface above a perfused fibroblast-lined microchannel. PM_10_ was applied to the epithelial surface for dose-exposure experiments.

## 2. MATERIALS & METHODS

### 2.1 VF-OOC Fabrication and Culture Systems

3D VF-OOC were developed using commercial BE-Transflow microfluidic devices (BEOnChip, Zaragoza, Spain). Each device had a culture microwell (volume 195 µL) positioned above a microchannel (4×1.5 mm, height 375 µm), separated by a porous polycarbonate membrane (thickness 10 µm, pore size 0.4 µm, pore density 1×10^5^ pores/cm^2^). Human primary VF fibroblasts (pVFF, passage 4 to 6), donated by the Thibeault Laboratory (University of Wisconsin-Madison), were embedded in type I collagen hydrogels (2.5 mg/mL, pH-adjusted to 7.2–7.4) at 0.5×10^6^ cells/mL. A total of 110 µL of pVFF-collagen gel was added per microwell and co-cultured with epithelial cells. Additional pVFF lined the microchannel to receive fluidic shear from continuous media perfusion at 40 µL/h using FAD media (1:3 DMEM/F12 with supplements), characterized by computational fluid dynamics (**Figure S1**), to approximate interstitial fluid flow velocities (0.1 to 4.0 µm/s) reported for soft tissues such as the VF mucosa [54-56].

Immortalized laryngeal epithelial cells (iLEC, ATCC CRL-3342, HuLa-PC) were seeded atop pVFF-collagen gels at 1×10^6^ cells/mL and co-cultured under submerged conditions for 3 days before an air-liquid interface (ALI) was established. iPSC-derived VF basal progenitor cells (iPSC line GM25256, Coriell Institute), differentiated following an established protocol [34], were seeded atop gels as a 10 µL drop and maintained in submerged culture for 2 to 4 days before ALI. Both iLEC and iPSC-derived cells were maintained at ALI with media perfusion through the microchannel for 10 days to allow epithelial stratification and maturation.

iLEC-based models (VF-OOC_iLEC_) were compared against three control conditions to isolate individual culture parameters. Namely, 3D transwell (static) cultures controlled for culture scaling effects. 2D chip (static) cultures (ChipShop, Jena, Germany) tested the influence of dimensionality. 3D chip (static) cultures isolated the combined effect of dynamic perfusion and fluidic shear, as these two parameters were applied together in the perfused chip configuration. iPSC-based models (VF-OOC_iPSC_) were compared against 3D transwell controls for verification. Conventional transwell cultures used the same cell types and co-culture configuration but without microfluidic perfusion. All cultures were performed as independent biological triplicates (N = 3). Detailed cell culture protocols and media compositions are provided in the Supplementary Methods.

### 2.2 Acute PM_10_ Exposure Experiments

Certified urban fine dust (ERM CZ100, European Institute for Reference Materials and Measurements) was used as the PM_10_ reference material [57]. ERM CZ100 is a well-characterized urban dust collected from a road tunnel with a certified mass median aerodynamic diameter below 10µm and known concentrations of polycyclic aromatic hydrocarbons, heavy metals, and inorganic ions. ERM CZ100 is routinely used in PM toxicity research [58-60]. At this initial developmental stage, using a standardized reference material was necessary to ensure that VF-OOC would generate reproducible and comparable results.

Determining PM_10_ concentrations for *in vitro* studies has been a methodological challenge. Ambient PM_10_ concentrations in air (∼59 µg/m^3^ daily average in the U.S. [61, 62]) translate to liquid-phase equivalents far below biologically detectable thresholds [63-65]. The concentrations used in this study (50 to 400 µg/mL) were thus selected to capture a range from low-level to intense exposure scenarios and are consistent with established *in vitro* PM toxicity protocols. Preliminary toxicity screening on pVFF and iLEC monocultures identified the onset of significant viability reduction at 100 µg/mL for pVFF (*p* < 0.05) and 400 µg/mL for iLEC (*p* < 0.05) (**Figure S2**). Based on these results, PM_10_ was applied for 24 hours across a dose range of 0 to 400 µg/mL for iLEC cultures and at 0 and 100 µg/mL for iPSC cultures, each in both VF-OOC and transwell formats. PM_10_ was suspended in FAD media from a stock solution (10 mg/mL) and applied directly to the epithelial surface in the microwell or transwell insert, mimicking natural particle deposition at the VF mucosal surface. All PM_10_ experiments included vehicle controls (0 µg/mL). All experiments were performed as independent biological triplicates (N = 3).

### 2.3 Microengineered VF Mucosal Tissue Characterization

Detailed characterization protocols are provided in the Supplementary Methods. In brief, the histology of microengineered VF mucosa was assessed by hematoxylin and eosin (H&E) staining of 10 µm cryosections, imaged on an Axiovert 3 Widefield Microscope (Zeiss) with a 40× objective. Immunocytochemistry was performed for cytokeratins, E-Cadherin, and vimentin (**Table 1**), imaged on an inverted Zeiss LSM710 confocal fluorescence microscope with a 20× objective and analyzed using Imaris 7.5.6 software (Bitplane). DAPI was used for counter-staining. Subcellular ultrastructure of the VF mucosal constructs was examined by transmission electron microscopy (TEM, Talos F200X STEM). TEM samples were sectioned at 70 nm thickness and stained with Reynolds lead citrate and uranyl acetate to increase image contrast.

**Table 1.** Reagents for immunocytochemistry and quantitative real-time PCR. AF = Alexa Fluorophore. *PM_10_ exposure experiments only.

| *Primary antibodies for immunocytochemistry* |  |  |  |  |  |  |  |
| --- | --- | --- | --- | --- | --- | --- | --- |
| Marker | Target | Host | Dilution | Catalogue No. | Fluorophore | Secondary Ab | Category |
| K5 | Cytokeratin 5 | Rabbit | 1:200 | Abcam ab193894 | AF488 | — | Stratified epithelium |
| K13 | Cytokeratin 13 | Rabbit | 1:200 | Abcam ab154346 | — | AF594 Goat anti-rabbit (1:500, ThermoFisher A-11037) | Stratified epithelium |
| K14 | Cytokeratin 14 | Mouse | 1:100 | Novus NBP2-34675AF532 | AF532 | — | Stratified epithelium |
| E-Cad | E-Cadherin | Rabbit | 1:100 | Abcam ab194982 | AF647 | — | Intercellular junction |
| Vimentin | Vimentin | Chicken | 1:300 | Abcam ab24525 | — | AF633 Goat anti-chicken (1:1000, ThermoFisher A-21103) | Intermediate filaments |

| DAPI | Nuclear DNA | — | 1:5000 | Abcam<br>ab228549 | — | — | Counter-staining |
| --- | --- | --- | --- | --- | --- | --- | --- |
| ***TaqMan assays for quantitative real-time PCR*** |  |  |  |  |  |  |  |
| Gene | Full Name | TaqMan Assay ID |  | Category |  |  |  |
| *KRT5* | Cytokeratin 5 | Hs00361185_m1 |  | Basal epithelium |  |  |  |
| *KRT13* | Cytokeratin 13 | Hs02558881_s1 |  | Suprabasal epithelium |  |  |  |
| *KRT14* | Cytokeratin 14 | Hs00265033_m1 |  | Basal epithelium |  |  |  |
| *TP63* | Tumor protein p63 | Hs00978340_m1 |  | Basal epithelium |  |  |  |
| *TJP1* | Tight junction protein 1 | Hs01551871_m1 |  | Tight junctions |  |  |  |
| *GJA1* | Gap junction alpha 1 | Hs00748445_s1 |  | Gap junctions |  |  |  |
| *MUC1* | Mucin 1 | Hs00159357_m1 |  | Mucus production |  |  |  |
| *HAS2* | Hyaluronan synthase 2 | Hs00193435_m1 |  | ECM synthesis |  |  |  |
| *HAS3* | Hyaluronan synthase 3 | Hs00193436_m1 |  | ECM synthesis |  |  |  |
| *PIEZO1* | Piezo-type mechanosensitive ion channel 1 | Hs00207230_m1 |  | Mechanotransduction |  |  |  |
| *TNF* | Tumor necrosis factor $\alpha$ | Hs00174128_m1 | | Cytokines* | | | |
| *IL6* | Interleukin 6 | Hs00174131_m1 |  | Cytokines* |  |  |  |
| *CXCL8 (IL-8)* | Interleukin 8 | Hs00174103_m1 |  | Cytokines* |  |  |  |
| *IL1A* | Interleukin 1α | Hs00174092_m1 | | Cytokines* | | | |
| *IL1B* | Interleukin 1β | Hs01555410_m1 | | Cytokines* | | | |
| *GAPDH* | GAPDH | Hs02786624_g1 |  | Housekeeping |  |  |  |

Gene transcriptions were measured by quantitative real-time PCR (qPCR) using TaqMan assays on a ViiA 7 system (ThermoFisher). Microtissue constructs were washed and submerged in collagenase type I (Gibco #17018029) until gel dissolution. Cells were collected and supernatant was aspirated. Cell pellets were stored at -80 °C until RNA extraction using the RNeasy Micro Kit (Qiagen #74004). Target genes included markers of epithelial stratification, tight and gap junctions, mucins, hyaluronan synthases, and the mechanosensitive ion channel *PIEZO1* (**Table 1**). Gene expression was reported relative to GAPDH using the 2^−ΔCt^ method.

For PM_10_ exposure experiments, gene expressions of inflammatory cytokines were further assayed (**Table 1**). For verification, secreted TNFα and IL-1β proteins were quantified by ELISA (Abcam) and normalized to total protein content via Bradford assay. Supernatant from both microchannels and microwells was collected and stored at -80°C until ELISA testing.

### 2.4 Computational Fluid Dynamics Analysis

Computational fluid dynamics (CFD) simulation was performed using Ansys Academic (Release 2023) to predict shear stress and velocity profiles in the VF-OOC microchannel at an experimental perfusion rate of 40 µL/h. Device geometry was obtained from CAD files provided by BEOnChip. The computational domain comprised 27,775 elements (element size 0.05 mm) with laminar flow assumptions and a no-slip wall boundary condition. Simulation parameters are detailed in the Supplementary Methods.

### 2.5 Statistical Analysis

Statistical assessment was performed using GraphPad Prism 10.3.1. One-way ANOVA or two-way between-subjects ANOVA were used as appropriate for each comparison, with assumptions verified prior to analysis. Dunnett, Sidak, or Holm-Sidak corrections were applied for multiple comparisons as appropriate. For iPSC dose-exposure data, pairwise comparisons across model types and across doses were treated as independent statistical evaluations and reported using Fisher’s LSD without multiple comparison corrections. Significance was set at p < 0.05.

## 3. RESULTS

### 3.1 Three-dimensional co-culture in microfluidic chips recapitulated VF mucosal architecture

To determine whether chip culture format and dimensionality affected VF mucosal tissue development, we compared 3D Transwell_iLEC_ (static), 2D VF-OOC_iLEC_ (static), and 3D VF-OOC_iLEC_ (static) cultures (**Figure 2a**). Histological staining revealed that 3D Transwell_iLEC_ and 3D VF-OOC_iLEC_ cultures both produced stratified squamous epithelia with distinct basal and suprabasal layers. Around 3-6 layers and 5-8 layers were observed in 3D Transwell_iLEC_ (static) and 3D VF-OOC_iLEC_ (static), respectively. In contrast, 2D VF-OOC_iLEC_ produced a thinner layer without clear stratification (**Figure 2b**). K5 and K14 expressions were confirmed in the basal epithelium across all conditions. E-Cadherin was detected in all conditions, with qualitatively stronger signal in the 3D Transwell_iLEC_ but minimal for 2D VF-OOC_iLEC_. Vimentin staining confirmed spindle-shaped fibroblasts within the collagen gel of both 3D formats.

**Figure 2.**
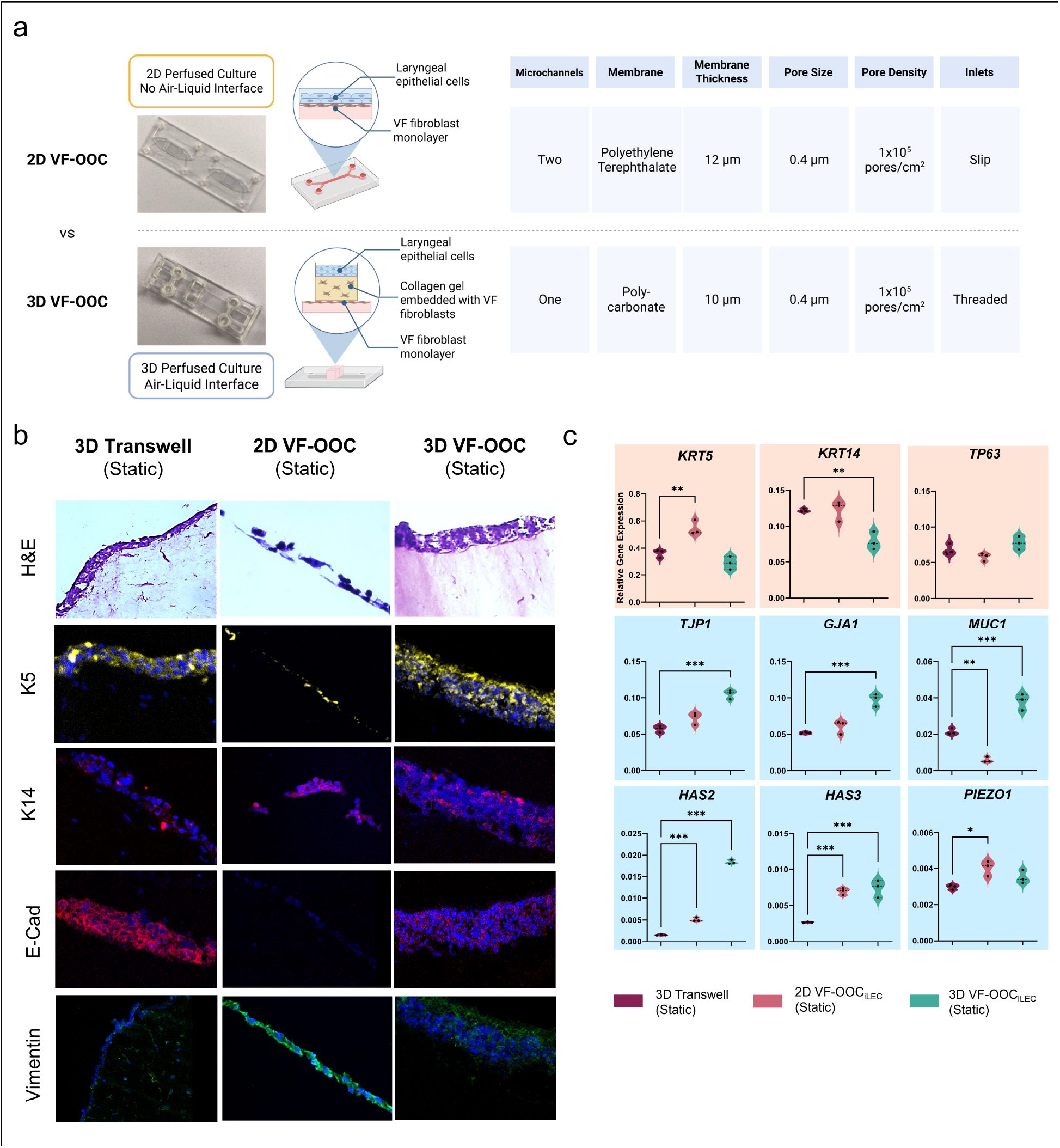
Effect of culture dimensionality and microfluidic scaling on VF mucosal tissue development. (a) Schematic comparison of 2D VF-OOC and 3D VF-OOC device configurations. (b) H&E and immunofluorescence (K5, K14, E-Cadherin, vimentin, DAPI) of 3D Transwell_iLEC_ (static), 2D VF-OOC_iLEC_ (static), and 3D VF-OOC_iLEC_ (static). H&E magnification 40×, scale bar = 15 μm. Immunostaining magnification 20×, scale bar = 30 μm. (c) Relative gene expression (calculated using 2^−ΔCt^ in comparison to GAPDH). N = 3. \**p*<0.05, \*\**p*<0.01, \*\*\**p*<0.001 by one-way ANOVA using Dunnett’s test for multiple comparisons.

Gene expression analysis revealed that chip cultures upregulated multiple gene categories relative to 3D Transwell_iLEC_ controls (**Figure 2c**). Barrier and junction genes *GJA1* (*p*<0.001) and *TJP1* (*p*<0.001) were significantly higher in the 3D VF-OOC_iLEC_ than Transwell_iLEC_. ECM remodeling genes *HAS2* and *HAS3* were significantly upregulated in both chip formats compared to Transwell_iLEC_ while the 3D VF-OOC_iLEC_ showed the highest expression (*p*<0.001 for all four comparisons). Among basal epithelial markers, *KRT5* was significantly higher in the 2D VF-OOC_iLEC_ (*p*<0.01) while the 3D VF-OOC_iLEC_ showed a comparable level to the Transwell_iLEC_ group. *KRT14* expression in the 3D VF-OOC_iLEC_ was lower than the Transwell_iLEC_ group (*p*<0.01). However, no significant difference was observed compared to the 2D VF-OOC_iLEC_ group. Although *KRT5* and *KRT14* gene expressions in the 3D VF-OOC_iLEC_ were not the highest among the three formats, K5 and K14 immunostaining (**Figure 2b**) showed a thicker epithelial interface with strong protein signal in the 3D VF-OOC_iLEC_, while the 2D VF-OOC_iLEC_ showed only a weak K5 signal at a thin interface.

All three culture formats showed a comparable level for the basal epithelial marker *TP63* expression. *PIEZO1* was significantly higher in the 2D VF-OOC_iLEC_ than Transwell_iLEC_ (*p*<0.05). The 3D VF-OOC_iLEC_ showed a comparable level to the 2D VF-OOC_iLEC_, with no significant difference from either the Transwell_iLEC_ or 2D VF-OOC_iLEC_ group. For mucin genes, *MUC1* was significantly lower in the 2D VF-OOC (*p*<0.01) and higher in the 3D VF-OOC (*p*<0.001) compared to Transwell_iLEC_. These results indicate that 3D VF-OOC_iLEC_ consistently upregulated barrier, mucin and ECM remodeling genes compared to 3D Transwell_iLEC_, while 2D VF-OOC_iLEC_ showed a more variable pattern with reduced mucin expression and elevated ECM remodeling gene expression.

### 3.2 Dynamic perfusion and fluidic shear promoted VF tissue maturation

We next evaluated the contribution of continuous perfusion and fluidic shear by comparing 3D Transwell_iLEC_ (static), 3D VF-OOC_iLEC_ (static), and 3D Dynamic VF-OOC_iLEC_ (perfused with pVFF-lined microchannel) cultures (**Figure 3**).

**Figure 3.**
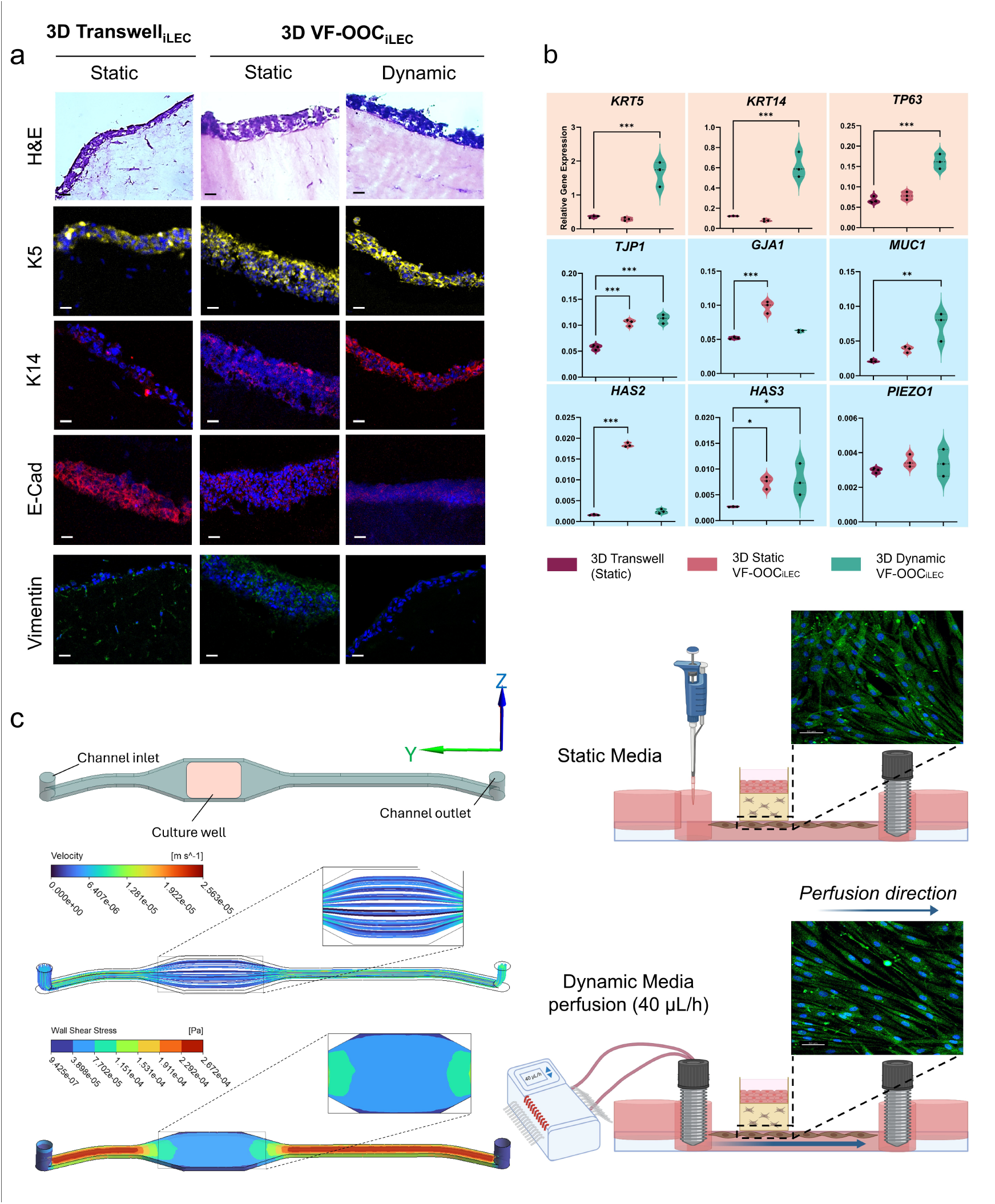
Effect of perfusion and fluidic shear on VF mucosal tissue maturation. (a) H&E and immunofluorescence of 3D Transwell_iLEC_ (static), 3D VF-OOC_iLEC_ (static), and 3D Dynamic VF-OOC_iLEC_. H&E magnification 40×, scale bar = 15 μm. Immunostaining magnification 20×, scale bar = 30 μm. (b) Relative gene expression (calculated using 2^−ΔCt^ in comparison to GAPDH). N = 3. \**p*<0.05, \*\**p*<0.01, \*\*\**p*<0.001 by one-way ANOVA using Dunnett’s test for multiple comparisons. (c) Left: CFD simulation of velocity and wall shear stress in the microchannel. Right: vimentin (green) and DAPI (blue) staining of pVFF in static versus dynamic conditions. Magnification 20×, scale bar = 50 μm.

Histology analyses showed that the 3D Dynamic VF-OOC_iLEC_ produced the most mature epithelium. The epithelium in the 3D Dynamic VF-OOC_iLEC_ was estimated ∼5-10 cell layers thick, compared to ∼5-8 layers in 3D VF-OOC_iLEC_ (static) and ∼3-6 layers in 3D Transwell_iLEC_. K5 and K14 signals appeared qualitatively stronger in the 3D Dynamic VF-OOC_iLEC_ compared to the other conditions (**Figure 3a**). E-Cadherin was detected across all conditions.

Gene expression profiling revealed that 3D dynamic chip cultures altered multiple gene categories relative to 3D Transwell_iLEC_ controls, with patterns varying by gene (**Figure 3b**). All three basal epithelial markers (*KRT5, KRT14, TP63*) showed the highest expression in the 3D Dynamic VF-OOC_iLEC_ and were significantly increased compared to 3D Transwell_iLEC_ (*p*<0.001). Barrier gene *TJP1* was significantly upregulated in both 3D chip formats versus 3D Transwell_iLEC_ (*p*<0.001), with the highest expression in the 3D Dynamic VF-OOC_iLEC_. Meanwhile, *GJA1* was significantly higher only in 3D Static VF-OOC_iLEC_ (*p*<0.001). Mucin gene *MUC1* was significantly higher in 3D Dynamic VF-OOC_iLEC_ (*p*<0.01 versus Transwell_iLEC_). ECM remodeling gene *HAS2* was markedly upregulated in 3D Static VF-OOC_iLEC_ (*p*<0.001 versus Transwell_iLEC_). In 3D Dynamic VF-OOC_iLEC_, *HAS2* showed a comparable level to Transwell_iLEC_. *HAS3* was significantly higher in both 3D chip formats compared to Transwell_iLEC_ where the highest expression was observed in the 3D Dynamic VF-OOC_iLEC_ (*p*<0.05 for both). *PIEZO1* showed comparable expression levels across the three conditions.

CFD simulations confirmed steady laminar flow with a parabolic velocity profile in the microchannel (**Figure 3c**). Wall shear stress varied along the channel. In the straight inlet and outlet portions, values ranged from ∼1.5-2.7 × 10^-4^ Pa. Beneath the culture well region where cells were cultured, values were lower, ranging from 3.9-7.7 × 10^-5^ Pa. Fibroblasts in the microchannel aligned in the direction of fluid flow under dynamic perfusion, as shown by vimentin and DAPI staining, while static controls showed random orientation (**Figure 3c**). Additional CFD parameters and results are provided in **Figure S1**.

### 3.3 VF-OOC reproduced subcellular ultrastructure of human VF mucosa

TEM of the 3D Dynamic VF-OOC_iLEC_ revealed subcellular features consistent with mature VF epithelium (**Figure 4a**). The superficial cell layers displayed flattened morphology, while deeper suprabasal cells were more cuboidal with visible cytoplasmic processes. Microvilli were present on the epithelial surface. Intercellular junctions (tight junctions, adherens junctions, desmosomes) were identified between adjacent epithelial cells (scale bar 500 nm). At the basal layer, a basement membrane with reticular fibers was visible (scale bar 1 µm). These features closely matched TEM images of adult human VF mucosa [66] (**Figure 4b**).

**Figure 4.**
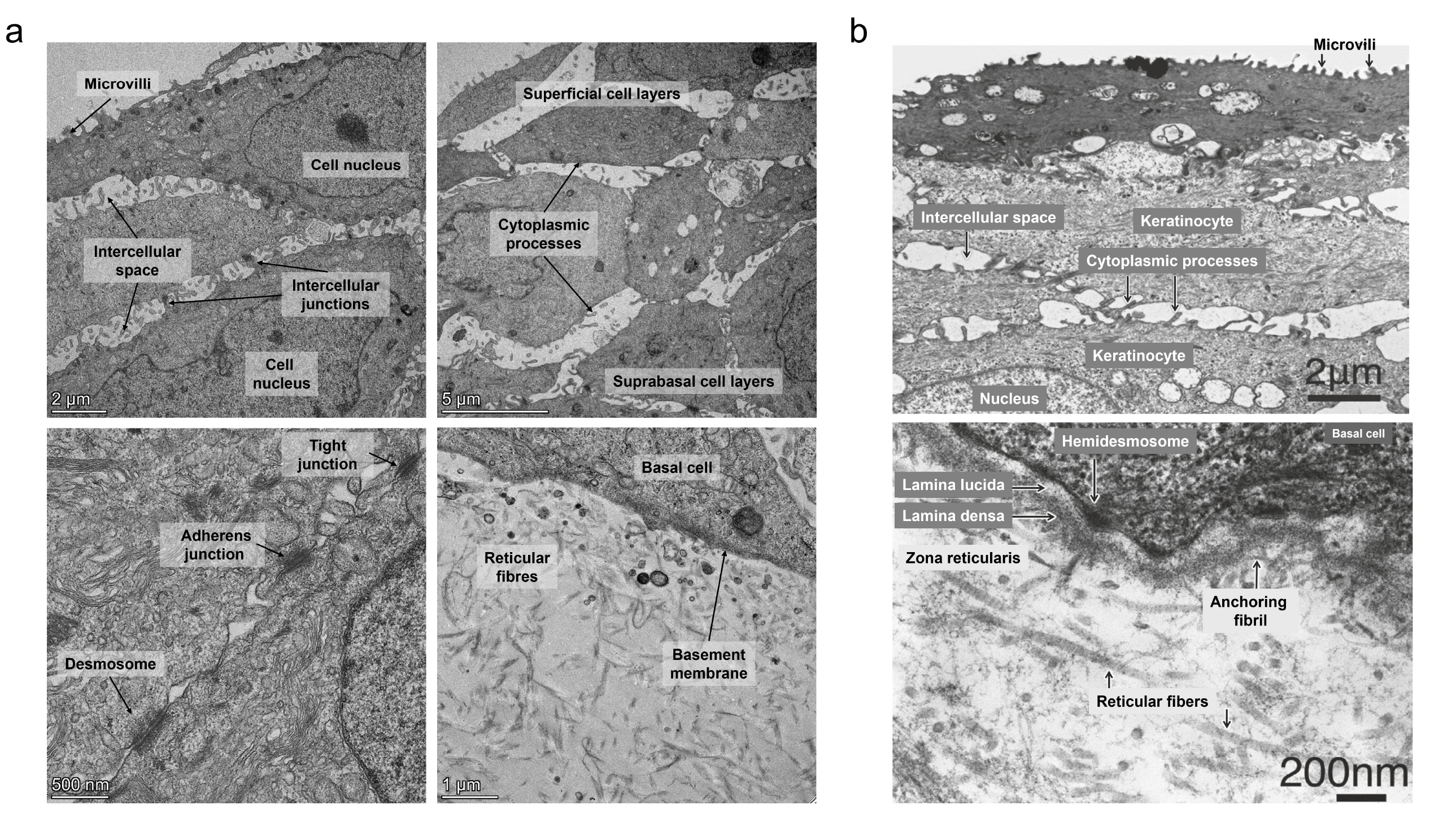
TEM of VF-OOC epithelium compared with native human VF mucosa. (a) 3D Dynamic VF-OOC_iLEC_. Top left: microvilli and intercellular junctions (scale bar 2 µm). Top right: suprabasal layers (scale bar 5 µm). Bottom left: tight junctions, adherens junctions, desmosomes (scale bar 500 nm). Bottom right: basement membrane with reticular fibers (scale bar 1 µm). (b) Adult human VF mucosa. Top: keratinocytes with microvilli (scale bar 2 µm). Bottom: basement membrane with lamina lucida, lamina densa, hemidesmosomes, anchoring fibrils (scale bar 200 nm). Panel (b) reprinted with permission from Sato K. (2018) Functional histoanatomy of the human larynx: Springer Singapore.

### 3.4 Acute PM_10_ challenge revealed platform-dependent mucosal responses

Transwell_iLEC_ and VF-OOC_iLEC_ were challenged with PM_10_ for 24 hours (**Figure 5**). Transmitted light microscopy showed PM_10_ particles accumulating at the epithelial surface in a dose-dependent manner (**Figure 5a**). Particles remained at or above the epithelial surface across all tested concentrations (0, 50, 100, and 400 µg/mL) in both platforms. K5, K14, E-Cadherin, and vimentin immunostaining were maintained across all doses. Epithelial structure and fibroblast morphology were preserved.

**Figure 5.**
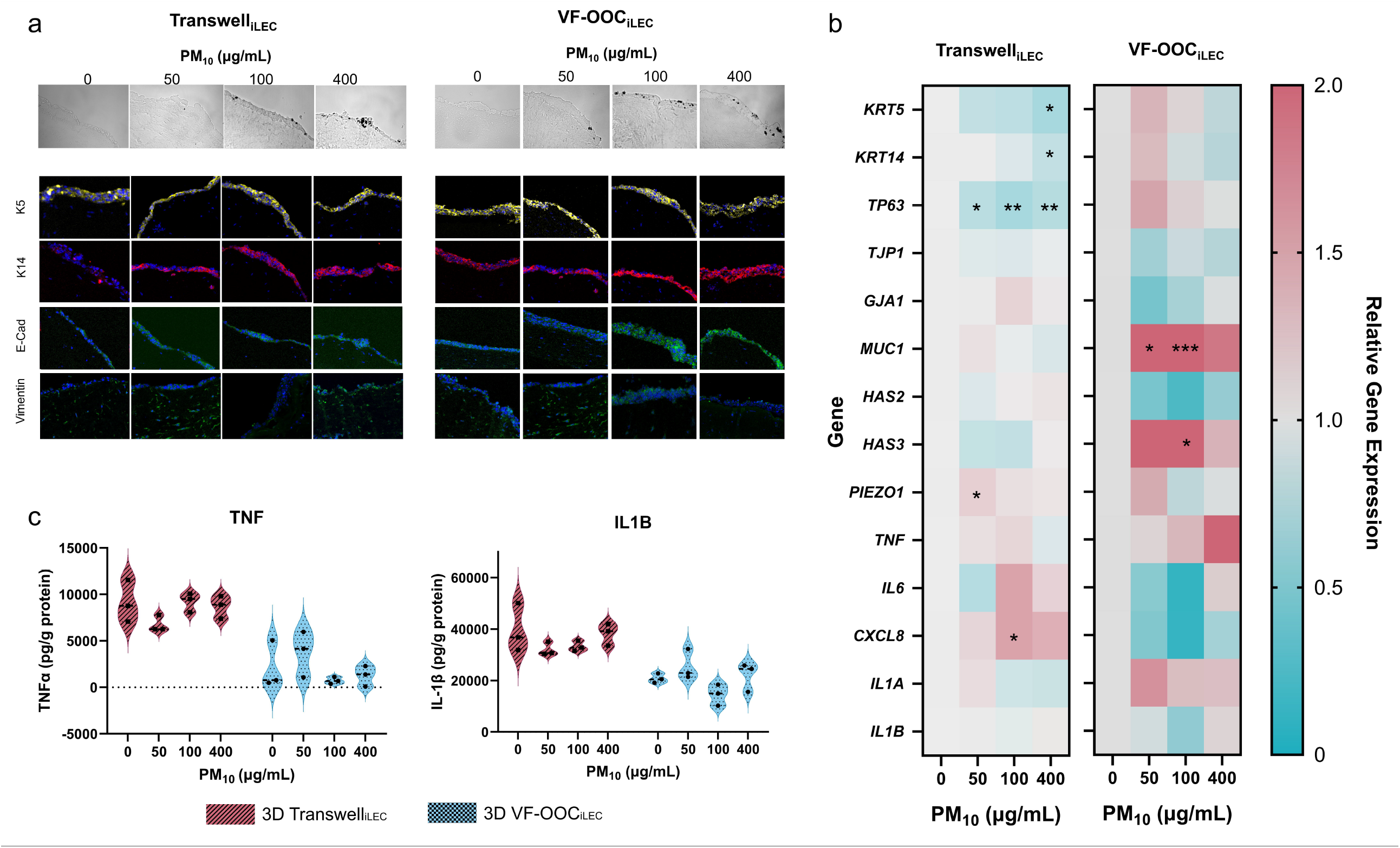
VF-OOC_iLEC_ response to acute PM_10_ challenge. (a) Transmitted light and immunofluorescence of Transwell_iLEC_ and VF-OOC_iLEC_ at 0, 50, 100, and 400 µg/mL PM_10_. Magnification 20×, scale bar = 30 μm. (b) Heatmap of relative gene expression, calculated using 2^−ΔCt^ in comparison to GAPDH and normalized to each gene’s 0 µg/mL control within each platform. Color represents gene expression relative to each gene’s 0 µg/mL control. Values exceeding 2.0 are displayed at the maximum scale. N = 3. \**p*<0.05, \*\**p*<0.01, \*\*\**p*<0.001 by one-way ANOVA versus 0 µg/mL control, using Sidak’s test for multiple comparisons. (c) Secreted TNFα and IL-1β (pg/g protein) by ELISA. N = 3. No significant dose-dependent differences were detected using one-way ANOVA.

Despite preserved staining for protein level expression (**Figure 5a**), transcriptional analysis revealed distinct response patterns between platforms (**Figure 5b**). Gene expression of 14 target genes (*KRT13* was excluded due to undetectable expression in iLEC) was normalized to their own 0 µg/mL controls within each platform. In Transwell_iLEC_, basal stratification markers were downregulated. *TP63* was progressively downregulated across PM_10_ doses (*p*<0.05 at 50 µg/mL, *p*<0.01 at 100 and 400 µg/mL). *KRT5* and *KRT14* were significantly downregulated at 400 µg/mL (*p*<0.05 for both). *PIEZO1* was significantly upregulated at 50 µg/mL (*p*<0.05) and *CXCL8* (*IL-8*) at 100 µg/mL (*p*<0.05). In VF-OOC_iLEC_, basal stratification markers showed no significant change, while mucin and ECM genes were upregulated. *MUC1* was significantly upregulated at 50 µg/mL (*p*<0.05) and 100 µg/mL (*p*<0.001), and *HAS3* at 100 µg/mL (*p*<0.05). The remaining genes showed no significant dose-dependent changes in either platform. Secreted cytokine levels showed no significant dose-dependent changes in either platform (**Figure 5c**). TNFα and IL-1β concentrations appeared generally higher in Transwell_iLEC_ than VF-OOC_iLEC_ across all doses, but no significant differences were detected between doses within either group.

In sum, transwell and chip VF mucosae maintained structural epithelial barrier integrity, with PM_10_ confined to the epithelial surface at all doses. However, the two platforms showed qualitatively different gene transcription patterns. In Transwell_iLEC_, three of the five dose-responsive genes were basal stratification markers, all of which were downregulated (*KRT5, KRT14, TP63*). In contrast, VF-OOC_iLEC_ showed only two dose-responsive genes, both upregulated barrier-supporting components (*MUC1* and *HAS3*), consistent with an adaptive mucosal response.

### 3.5 iPSC-derived VF-OOC detected cytokeratin 13 upregulation after PM_10_ exposure

To test whether the platform-dependent response pattern observed with iLEC also occurred in a more terminally differentiated epithelial source, we developed a VF-OOC using iPSC-derived VF basal progenitor cells for the acute PM_10_ exposure experiment (**Figure 6**).

**Figure 6.**
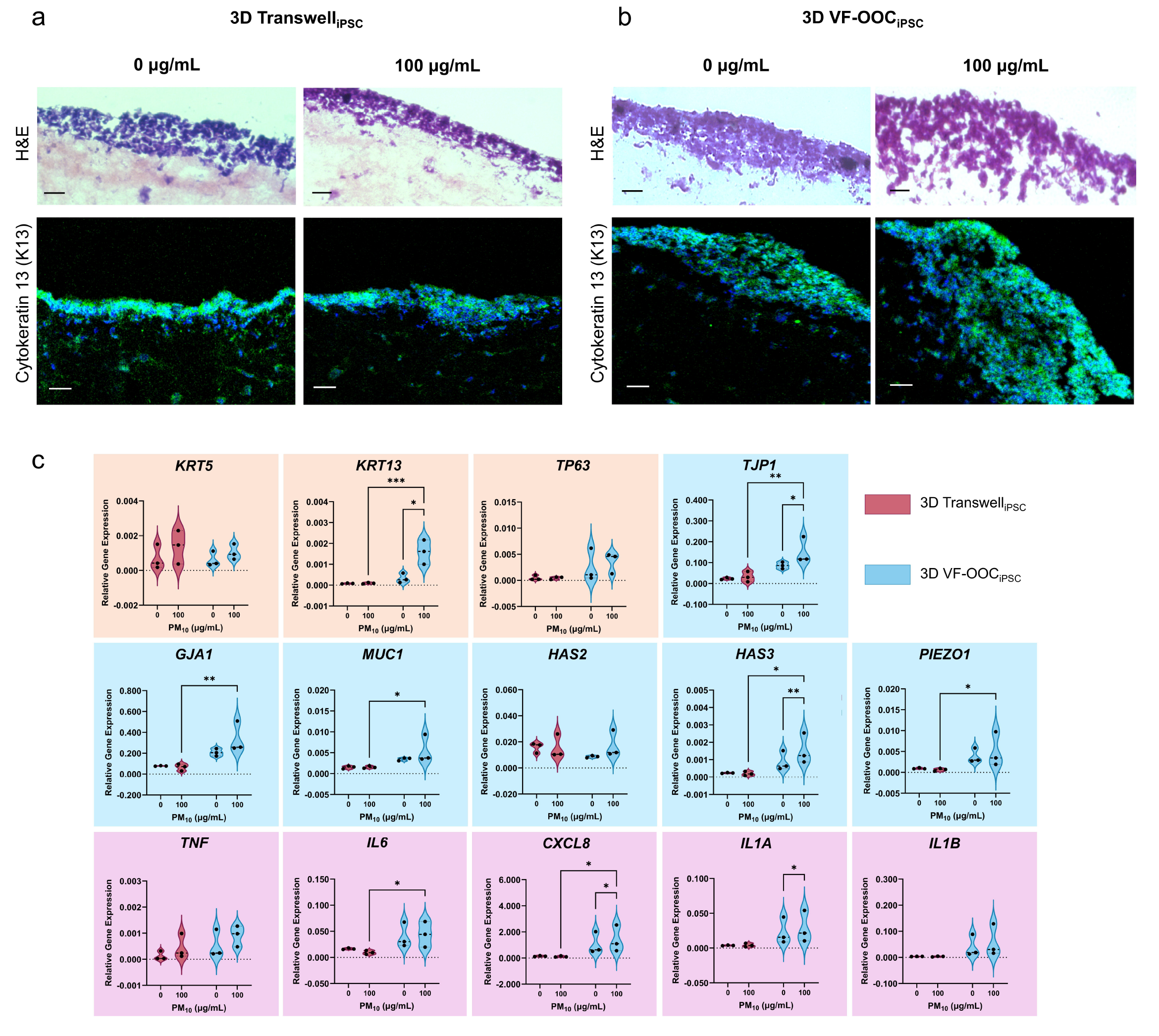
iPSC-derived VF-OOC response to PM_10_ exposure. (a) Transwell_iPSC_ and (b) VF-OOC_iPSC_ H&E and K13 immunofluorescence at 0 and 100 µg/mL PM_10_. H&E magnification 40×, scale bar = 15 μm. Immunostaining magnification 20×, scale bar = 30 μm. (c) Relative gene expression (calculated using 2^−ΔCt^ in comparison to GAPDH). K14 was below detection and excluded. N = 3. \**p*<0.05, \*\**p*<0.01, \*\*\**p*<0.001 by two-way ANOVA. Pairwise comparisons across model types and across doses were treated as independent statistical evaluations. Therefore, the reported *p*-values derive from Fisher’s LSD without multiple comparison corrections.

Overall, VF-OOC_iPSC_ produced a visually thicker, more densely stratified epithelium than Transwell_iPSC_ at baseline (**Figure 6a, b**, 0 µg/mL). After PM_10_ exposure, the VF-OOC_iPSC_ epithelium appeared slightly less compact. Also, VF-OOC_iPSC_ displayed stronger, more widespread K13 signals throughout the epithelium than those of Transwell_iPSC_. K13 was absent in all iLEC-based models (data not shown), consistent with the known limited differentiation capacity of the immortalized cell line.

Gene expression analysis revealed both dose-dependent and platform-dependent differences (**Figure 6c**). Within the VF-OOC_iPSC_ platform, five genes associated with epithelial differentiation (*KRT13*), barrier function (*TJP1*), ECM remodeling (*HAS3*), and inflammation (*CXCL8, IL1A*) were significantly upregulated by the PM_10_ dose challenge (0 versus 100 µg/mL). In comparing VF-OOC_iPSC_ to Transwell_iPSC_ at 100 µg/mL, *KRT13* (*p*< 0.001), *TJP1* (*p*<0.01), *HAS3* (*p*<0.05), and *CXCL8* (*p*<0.05) were significantly higher in VF-OOC_iPSC_. Other genes including *GJA1* (*p*<0.01), *MUC1* (*p*<0.05), *PIEZO1* (*p*<0.05), and *IL6* (*p*<0.05) were also significantly higher in VF-OOC_iPSC_ at 100 µg/mL. These results are consistent with the platform-dependent effects observed in the iLEC experiments (**Figures 2 and 3**).

Two genes were dose-responsive in both iPSC and iLEC formats. *HAS3* was upregulated in both VF-OOC_iPSC_ and VF-OOC_iLEC_, and *CXCL8* was upregulated in both VF-OOC_iPSC_ and Transwell_iLEC_ (**Figure 5b**). *KRT5, TP63, HAS2, TNF*, and *IL1B* showed no significant differences in any comparison. *KRT14* was below the qPCR detection limit in most iPSC-derived samples and was excluded from the analysis. Key findings from both VF-OOC development and PM_10_ dose-exposure experiments are summarized in Table 2.

**Table 2.** Summary of Key Findings.

|  | Transwell<sub>iLEC</sub> | VF-OOC<sub>iLEC</sub> (dynamic) | Transwell<sub>iPSC</sub> | VF-OOC<sub>iPSC</sub> (dynamic) |
| --- | --- | --- | --- | --- |
| **Throughput** | High | Low | High | Low |
| **Cost** | Low | High | Low | High |
| **Cell requirement** | >50,000 | <50,000 | >50,000 | <50,000 |
| **Media volume** | High | Low | High | Low |
| **Perfusion** | No | Yes | No | Yes |
| **Epithelium thickness** | ~3-6 layers | ~5-10 layers | ~3-6 layers | ~5-10+ layers |
| **Epithelium** | Stratified, basal and suprabasal layers | Most organized under dynamic perfusion with visible tight junctions, adherens junctions, desmosomes, reticular fibers | Terminally differentiated, flattened outer layer | Terminally differentiated, thicker than Transwell |
| **Cytokeratin expression** | K5, K14 (K13 undetectable) | K5, K14 (K13 undetectable) | K13, K5 detected at gene level, K14 undetectable | K13, K5 detected at gene level, K14 undetectable |
| **Barrier genes** | Reference | *TJP1* significantly higher, *GJA1* not significant | Reference | *TJP1* significantly higher, *GJA1* significantly higher |
| **Mucin genes** | Reference | *MUC1* significantly | Reference | *MUC1* significantly |
|  |  | higher |  | higher |
| **ECM genes** | Reference | *HAS2* not significant,<br>*HAS3* significantly higher | Reference | *HAS2* not significant,<br>*HAS3* significantly higher |
| **PM<sub>10</sub> dose-responsive genes** | *TP63*, *KRT5*, *KRT14*,<br>*PIEZO1*, *CXCL8* | *MUC1*, *HAS3* | None | *KRT13*, *TJP1*, *HAS3*,<br>*CXCL8*, *IL1A* |

## 4. DISCUSSION

This study presents a chip-based coculture model of human VF mucosa. We applied PM_10_ to this biomimetic platform to match the inhaled size fraction predicted to deposit at this anatomical site. Most airway on-a-chip work has targeted bronchial and alveolar epithelium [43, 49, 67-69]. VF mucosa has stratified squamous epithelium overlying a fibroblast-rich lamina propria. Bronchial pseudostratified ciliated platforms do not reproduce this architecture. Most existing *in vitro* VF mucosa models are mainly static monocultures that use either epithelial cells at ALI [32] or fibroblasts in submerged culture [28, 29]. The closest *prior* culture platform was a 3D iPSC-derived VF co-culture with stratified squamous architecture but lacked fluidic perfusion [34].

In this study, VF-OOC reproduced key epithelial and subcellular features of adult human VF and preserved this architecture under PM_10_ challenge. The mucin gene (*MUC1*) and ECM remodeling genes (*HAS2* and *HAS3*) were also upregulated in the 3D chip relative to 3D Transwell controls. In iLEC cultures, basal stratification markers (*KRT5, KRT14, TP63*) showed the highest expression in the dynamic chip. Intercellular junctional complexes and a defined basement membrane characteristic of adult human VF further confirmed this biological representation (**Figure 4**) [66, 70]. Validation across both iLEC-and iPSC-derived VF epithelium showed that the chip accommodates two distinct cell sources. Under PM_10_ challenge, particles deposited on the epithelial surface and did not penetrate beyond the basement membrane in either transwell or chip cultures across all PM_10_ doses. Tissue architecture remained intact as K5, K14, E-Cadherin, and vimentin stains were maintained.

Transwell and VF-OOC formats produced distinct transcriptional patterns under PM_10_ challenge. In Transwell_iLEC_, *TP63* was downregulated at all three doses, *KRT5* and *KRT14* were downregulated at 400 µg/mL, *CXCL8* was induced at 100 µg/mL, and *PIEZO1* was induced at 50 µg/mL. K5 and K14 protein staining remained intact across doses, consistent with the slower turnover of stable cytokeratins within the 24-hr acute window. VF-OOC_iLEC_ showed upregulation of barrier components *MUC1* (at 50 and 100 µg/mL) and *HAS3* (at 100 µg/mL), suggesting an adaptive mucosal response. Secreted TNFα and IL-1β showed no dose-dependent change in either format. Consistently, no significant differences were observed in *TNF* and *IL1B* gene expression. Baseline cytokine concentrations were nonetheless higher in Transwell_iLEC_ than in VF-OOC_iLEC_, likely reflecting accumulation in static media versus continuous clearance under chip perfusion.

The acute mucosal response to PM_10_ differs from soluble inhaled exposures because particles are mostly confined above the VF stratified epithelium and basement membrane. In our cultures, PM_10_ remained at the epithelial surface across all doses, and the VF-OOC_iLEC_ response shifted toward barrier reinforcement (*MUC1, HAS3* upregulation) rather than the proinflammatory gene induction reported when soluble cigarette smoke extract was applied to a 3D VF mucosal model and diffused through the epithelium [34]. Prior 2D pVFF monocultures, in which particles directly contacted fibroblasts, showed NLRP3-mediated pyroptosis at 20 µg/mL [29] and MAPK/NF-κB activation at higher doses [28]. This inflammatory response was not reproduced in our coculture models, even at doses up to 400 µg/mL, because the stratified epithelium in the 3D coculture shielded the fibroblast compartment from direct particle contact, as occurs *in vivo*.

While the VF-OOC_iLEC_ model showed barrier reinforcement to PM_10_, the VF-OOC_iPSC_ model at 100 µg/mL showed induction of genes for suprabasal differentiation (*KRT13*), tight junctions (*TJP1*), ECM remodeling (*HAS3*), and inflammation (*CXCL8, IL1A*). Of these five iPSC-induced genes, *HAS3* induction was also observed in VF-OOC_iLEC_ and *CXCL8* in Transwell_iLEC_. *KRT13, TJP1*, and *IL1A* responses were specifically observed in VF-OOC_iPSC_. In native human VF, cytokeratin 13 localizes to suprabasal cell layers [66]. In contrast, the commercial iLEC line in this study was from the posterior glottis with limited terminal differentiation capacity [33]. This basal bias might explain the absence of *KRT13* in iLEC cultures. Conversely, iPSC-derived VF basal progenitors were at an earlier and less stable differentiation state with a suprabasal leaning. Because basal cytokeratins require stable basal commitment, this progenitor state could account for the variable *KRT5* and undetectable *KRT14* in iPSC cultures. That said, the concurrent upregulation of *CXCL8* and *IL1A* in VF-OOC_iPSC_ matched inflammatory profiles of chronic laryngitis and PM-associated upper airway disease [19]. For toxicological applications requiring sensitivity to differentiation-related endpoints, iPSC-derived VF epithelium may be preferable.

Immune cells were not included in this first version of VF-OOC, which constrains its ability to model immune-mediated responses. PM_10_ exposure has been associated with delayed presentation of laryngitis admissions [26], a time-lag that suggests downstream immune activation rather than direct epithelial injury. The observation of limited cytokine response in this study is consistent with a human skin model in which PM-induced proinflammatory cytokine release was detected only in the presence of resident Langerhans cells [71]. Future VF-OOC iterations incorporating macrophages or dendritic cells will allow direct testing of immune-mediated mechanisms in pollutant-associated upper airway disease. That said, adding immune cells to the VF-OOC is non-trivial because shared culture media must preserve the phenotype and viability of all cell types, a known technical challenge in multi-cell-type co-cultures [34, 72].

Although ultrastructural TEM evidence supported barrier integrity, quantitative permeability was not measured. TEER is validated for conventional culture inserts instead of microfluidic chips. A functional permeability assay is a development need for chip-specific configuration. Also, *KRT13* was the only suprabasal marker measured in VF-OOC_iPSC_. A broader differentiation panel is needed to confirm this cellular shift. PM_10_ was applied as a liquid suspension to submerged cultures after ALI maturation, an exposure route typical of VF studies that model inhaled agents through soluble surrogates such as cigarette smoke extract [34, 73]. A submerged format was chosen because aerosol exposure of microfluidic chips is difficult, as current delivery systems target transwell plate formats [74]. Adapting aerosol delivery and extending exposure to chronic timescales are future directions.

Here, we presented an OOC model of human VF mucosa and applied it to acute PM_10_ exposure. Following this exposure, VF-OOC demonstrated an adaptive mucosal response, with upregulation of barrier-supporting components *MUC1* and *HAS3* and stable inflammatory cytokine signaling. Animal models cannot address laryngeal pathophysiology directly due to different anatomical and molecular features. Further development of VF-OOC will advance the NAM toolset in laryngology research. This platform extends respiratory toxicology from lungs to upper airways and can be applied to other inhaled exposures such as wildfire smoke, occupational dust and e-cigarette aerosols. Ultimately, the VF-OOC supports mechanistic studies linking these exposures to laryngeal conditions such as chronic laryngitis and cancer.

## Author Contributions

P.T.C.: Conceptualization, Methodology, Investigation, Writing-Review.

M.M: Software, Formal analysis, Data curation, Validation, Visualization, Writing-Original Draft, Writing-Review & Editing.

Y.L: Formal analysis, Validation, Data curation, Writing-Original Draft, Writing-Review & Editing.

V.L.: Conceptualization, Methodology, Writing-Review & Editing.

S.T.: Visualization, Validation, Writing-Review & Editing.

L.M.: Formal analysis, Validation, Data curation, Writing-Review.

X.L.: Conceptualization, Writing-Review & Editing.

G.M.: Formal analysis, Validation, Data curation, Writing-Review.

C.X.-Q.C.: Methodology, Writing-Review & Editing.

T.D.: Methodology, Writing-Review & Editing.

S.L.T.: Conceptualization, Writing-Review & Editing.

N.Y.K.L.-J.: Conceptualization, Validation, Visualization, Resources, Writing-Original Draft, Writing-Review & Editing, Supervision, Funding acquisition, and Project administration.

## Acknowledgements

We would like to thank McGill Facilities for Advanced BioImaging (ABIF) and Imaging and Molecular Biology Platform (IMBP, Dr. Nicolas Audet) for technical support.

## Data Availability Statement

All data and code that support the findings of this study are available from the corresponding author upon reasonable request.

