## Supplementary Methods for "A Human Vocal Fold Organ-On-Chip for Studying Platform-Dependent Mucosal Responses to Particulate Matter"

### *Development of a Vocal Fold Organ-on-Chip for Modeling Upper Airway Mucosal Responses to Particulate Matter*

#### S1. Cell Culture and Maintenance

**S1.1 Human Primary Vocal Fold Fibroblasts (pVFF)**

pVFF were maintained in DMEM (Sigma #D6429) supplemented with 10% (v/v) FBS (ThermoFisher #12484028), 1% (v/v) penicillin-streptomycin (ThermoFisher #15140122), and 1% (v/v) non-essential amino acids (ThermoFisher #11140050).

**S1.2 Immortalized Laryngeal Epithelial Cells (iLEC)**

iLEC (ATCC CRL-3342, HuLa-PC) were maintained in dermal cell basal medium (ATCC #PCS-200-030) supplemented with a keratinocyte growth kit (ATCC #PCS-200-040) per manufacturer recommendations. During experiments, iLEC were cultured in stratified DMEM for 72 hours prior to seeding. To prepare stratified DMEM, basal DMEM was first made by supplementing DMEM/F12 (ThermoFisher #10565018) with 2% B27 (ThermoFisher #17504044), 1% N2 (ThermoFisher #17502048), and 1% penicillin-streptomycin. Stratified DMEM was then prepared by adding 0.4 µg/mL hydrocortisone (Stemcell Technologies #07925), 8.4 ng/mL cholera toxin (Sigma #C9903), 5 µg/mL insulin (Sigma #91077C), 24 µg/mL adenine (Sigma #A8626), and 20 ng/mL EGF (Stemcell Technologies #78006).

**S1.3 iPSC-derived VF Basal Progenitor Cells**

Protocols from the Montreal Neurological Institute were followed for maintaining iPSC in an undifferentiated state. Matrigel coating was prepared on ice by adding 100 µL of Matrigel (100×) (Corning #354277) to 10 mL of DMEM/F12 with Antibiotic-Antimycotic (1×) (ThermoFisher #15240096). Coated 60 mm petri dishes (Eppendorf #0030701119) were incubated for 1 hour at 37 °C. iPSC (GM25256, Coriell Institute) in complete mTeSR Plus (Stemcell Technologies #100-0276) supplemented with 10 μM ROCK inhibitor (Stemcell Technologies #72302) were seeded in coated dishes and incubated for 20 hours, at which point complete mTeSR Plus was added. Media was changed daily, and iPSC were routinely passaged using ReLeSR (Stemcell Technologies #05872) at 60–70% confluency.

At 80% confluency, iPSC were subjected to the differentiation protocol [1]. On day 1, definitive endoderm differentiation was initiated with RPMI media with Glutamax (ThermoFisher #61870-036), 100 ng/mL Activin A (Stemcell Technologies #78001), 25 ng/mL Wnt 3a (R&D Systems #5036-WN-010), and 10 μM ROCK inhibitor for 24 hours. On day 2, the media was changed to RPMI with Glutamax, 100 ng/mL Activin A, and 0.2% FBS. Cells were incubated for 48 hours with media replenished after 24 hours.

On day 4, anterior foregut endoderm differentiation commenced with basal DMEM, 200 ng/mL noggin (Stemcell Technologies #78060), 10 μM SB431542 (Stemcell Technologies #72232), 50 µg/mL ascorbic acid (Stemcell Technologies #72132), and 0.4 mM 1-thioglycerol (Sigma #M6165). Cells were incubated for 96 hours with media refreshed daily. On day 9, VF basal progenitor differentiation was induced with basal DMEM, 50 µg/mL ascorbic acid, 250 ng/mL FGF-2 (Stemcell Technologies #78003), 100 ng/mL FGF-7 (Stemcell Technologies #78046), and 100 ng/mL FGF-10 (Stemcell Technologies #78037) for 48 hours. On day 10, culture quality was assessed by brightfield microscopy. Proliferation, high confluency, and limited cell stacking indicated successful progenitor differentiation.

**S1.4 FAD Media Preparation**

FAD media was prepared from DMEM medium-high glucose (Sigma #D6429) and Ham’s F12 (ThermoFisher #11765054) mixed in a 1:3 ratio, supplemented with 2.5% FBS, 1% penicillin-streptomycin, 0.4 µg/mL hydrocortisone, 8.4 ng/mL cholera toxin, 5 µg/mL insulin (Sigma #91077C), 24 µg/mL adenine (Sigma #A8626), and 10 ng/mL EGF.

**S1.5 Conditional FAD Media Preparation**

For iPSC-based cultures, conditional FAD media was prepared by applying FAD media to a stock pVFF culture at 60–70% confluency and incubating overnight. The conditioned media was collected, sterile-filtered, aliquoted, and frozen at −20 °C until required. Aliquots were mixed with fresh FAD media in a 30:70 ratio to generate conditional FAD media.

#### S2. Culture System Setup

**S2.1 VF-OOCᵢLEC Setup**

The microchannel of the BE-Transflow device was coated with collagen (50 µg/mL) prior to seeding pVFFs as a monolayer and incubating for 24 hours. Collagen gels (2.5 mg/mL) (ThermoFisher #A10483-01) were fabricated on ice and adjusted to pH 7.2–7.4 using 1 N NaOH (ThermoFisher #124260010). pVFFs in FBS were added to produce a final concentration of 0.5×10^6^ cells/mL. 110 µL of pVFF-collagen gel mix was added to the culture well and incubated for 75 minutes to initiate gel formation. Gels were detached from the membrane and basal DMEM was added to the well and channel. Devices were incubated for 24 hours to facilitate gel contraction.

After 24 hours, iLEC were seeded atop gels at a concentration of 1×10^6^ cells/mL and incubated to allow cell attachment. Cells were co-cultured under submerged conditions for three days before an ALI was established by removing media from the well. The microfluidic device was then connected to a peristaltic pump and FAD media was perfused through the microchannel at 40 µL/h for 10 days.

Static controls were prepared as above but were not connected to a peristaltic pump, with daily media exchanges performed via pipetting. All variations of VF-OOC_iLEC_ culture were performed as independent biological triplicates.

**S2.2 TranswellᵢLEC Setup**

Transwell culture systems used both pVFF and iLEC grown on conventional transwell inserts. Collagen gels were prepared by combining 6.7 mL collagen I (ThermoFisher #A10483-01), 1 mL 10× MEM (ThermoFisher #11430030), and 2.13 mL of distilled water (ThermoFisher #15230170) on ice. Collagen gel pH was adjusted to 7.2–7.4 using 1 N NaOH and verified with pH strips.

pVFFs were pelleted and resuspended in ice-cold FBS at 5×10^6^ cells/mL. 1 mL of cell suspension was added to the collagen gel on ice to generate a final concentration of 0.5×10^6^ cells/mL. 110 µL of gel-cell mix was dispensed into the insert of a 24-well transwell plate and incubated at 37 °C for 75 minutes to initiate gel formation. Gels were gently detached from the membrane and 600 µL of basal DMEM added to each well alongside 100 µL to each insert. The plate was incubated for a further 24 hours to allow gel contraction.

iLECs were harvested, counted, pelleted, and resuspended in stratified DMEM at 1×10^6^ cells/mL. All media was removed from the transwell plate and 100 µL of iLECs added atop each collagen gel. 600 µL of stratified DMEM was also added to the well. Media was changed every 48 hours. Cells were co-cultured in submerged conditions for 3 days to allow attachment and proliferation. Three days post-iLEC seeding, an ALI was established by removing media from the insert. The media in the well was changed to FAD. Well media was changed every 48 hours. Cells were cultivated at the ALI for a further 10 days. All transwell_iLEC_ cultures were performed as independent biological triplicates.

**S2.3 VF-OOCᵢPSC Setup**

VF-OOC_iPSC_ used the same initial microchannel coating and collagen-pVFF gel preparation as VF-OOC_iLEC_, except for the epithelial cell source. After 24 hours for gel contraction, VF basal progenitors were harvested via TrypLE Express, resuspended in stratified DMEM supplemented with FGFs, and seeded atop gels as a 10 µL drop. After progenitor seeding, devices were incubated for 2 hours to allow attachment. Stratified DMEM with FGFs was then added to the well and microchannel. Media was changed daily.

Two days after VF basal progenitor seeding, all media was changed to conditional FAD media supplemented with FGFs. Submerged culture was maintained for 2–4 days to permit cell proliferation to cover the gel. After epithelial confluency was reached, an ALI was established by removing media from the well. The microfluidic device was then connected to a peristaltic pump and FAD media was perfused through the channel at 40 µL/h. Cells were cultivated in this perfused ALI setup for a further 10 days. All VF-OOC_iPSC_ cultures were performed as independent biological triplicates.

**S2.4 TranswellᵢPSC Setup**

Transwell_iPSC_ systems used the same initial collagen-pVFF gel preparation as VF-OOC_iPSC_. VF basal progenitors were harvested using TrypLE Express, resuspended in 200 µL of stratified DMEM supplemented with FGFs, and seeded atop gels as a 10 µL drop following the protocol of Lungova et al. [1]. After progenitor seeding, transwell plates were incubated for 2 hours to allow attachment. Stratified DMEM with FGFs was added to the insert and well. Media was changed daily. Cells were cultured in submerged conditions for a further 48 hours. Two days after seeding, all media was changed to conditional FAD media supplemented with FGFs. Submerged culture was maintained for 2–4 days to allow cell proliferation. After epithelial confluency was reached, an ALI was established by removing media from the insert. Cells were maintained at an ALI for a further 10 days. All transwell_iPSC_ cultures were performed as independent biological triplicates.

**S2.5 2D VF-OOC Setup**

The 2D VF-OOC used Cross-Flow microfluidic devices (Microfluidic ChipShop GmbH, Jena, Germany, #10001555) consisting of two parallel chambers (upper 220 µL, lower 120 µL) separated by a porous polyethylene terephthalate membrane (thickness 12 µm, pore size 0.4 µm, pore density 1×10^5^ pores/cm^2^). Channels were filled with DMEM and incubated overnight to equilibrate the device. Channels were then coated with collagen by pipetting a solution of 50 µL collagen in 3 mL DMEM into each channel and incubating for 1 hour at 37 °C. After washing, channels were refilled with DMEM until cell seeding.

The lower channel was filled with pVFF resuspended in basal DMEM at 0.5×10^6^ cells/mL. The chip was inverted and incubated overnight to permit attachment, then incubated for a further 48 hours with media changed at 24 hours. iLEC were resuspended in stratified DMEM at 1×10^6^ cells/mL and applied to the upper channel 48 hours after pVFF seeding. Media in the lower channel was also changed to stratified DMEM. Media in both channels was changed every 24 hours via pipetting. 2D VF-OOC co-cultures were maintained in submerged conditions for 13 days, without an air-liquid interface or collagen gel embedding. All 2D VF-OOC cultures were performed as independent biological triplicates.

#### S3. Computational Fluid Dynamics

Device geometry was obtained from CAD files provided by BEOnChip and imported into Ansys Academic (Release 2023). The computational domain comprised 27,775 elements (element size 0.5 mm) with laminar flow assumptions and a no-slip wall boundary condition. Material properties of the channel and fluid were specified using those of plastic (ρ = 1.2 g/cm^3^) and DMEM supplemented with 10% FBS (ρ = 1.009 g/cm^3^) respectively. The input velocity was derived from the experimental flow rate as follows.

*v* = *Q* / *A*

where *Q* is the volumetric flow rate, *A* is the cross-sectional area, and *v* is the average velocity. The calculation was performed for 750 iterations and scaled residuals (continuity and velocity) were monitored to ensure convergence. Velocity and wall shear stress profiles are shown in Figure S2.

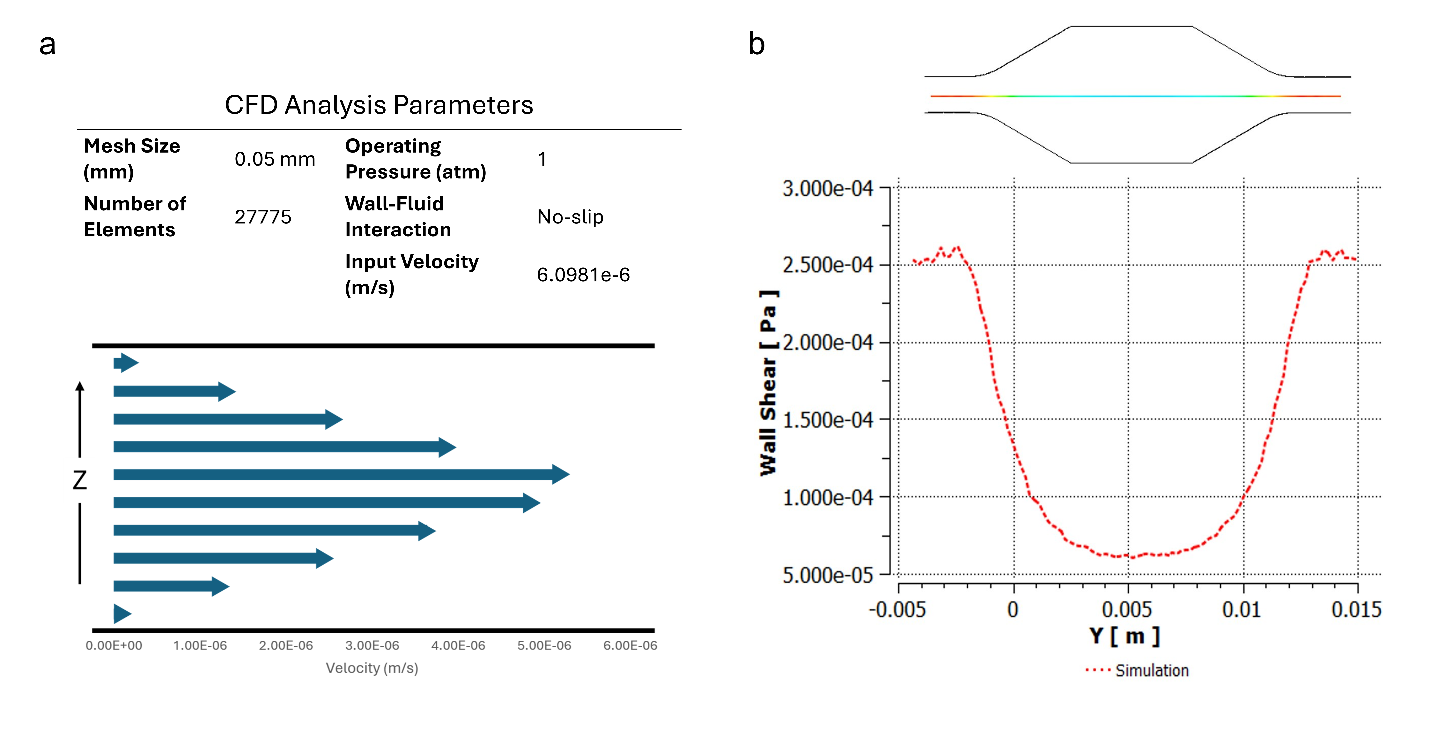
**Figure S1. CFD simulation parameters and additional shear stress profiles.** (a) Parabolic velocity profile across microchannel height, consistent with laminar flow. (b) Wall shear stress along channel length, with low, uniform shear underneath the culture well. See table for mesh and flow parameters.

#### S4. PM₁₀ Exposure Experiments

Certified urban fine dust ERM CZ100 (European Institute for Reference Materials and Measurements) was used as the PM_10_ reference material [2]. Working concentrations were prepared from a stock solution (10 mg/mL) by adding 100 mg PM_10_ to 10 mL of FAD media and vortexing for 5 minutes. For preliminary monoculture toxicity screening, pVFF and iLEC were seeded separately in 24-well cell culture plates (Eppendorf #0030741021) at 0.5×10^5^ cells/mL. After cultures reached confluency, they were exposed to PM_10_ at 0, 25, 50, 100, 200, 400, 800, and 1600 µg/mL for 24 hours. Following exposure, monoculture viability was evaluated by LIVE/DEAD cytotoxicity assay (Invitrogen #1932445) (Figure S1).

For co-culture exposure experiments, PM_10_ in FAD media was applied directly to the epithelial surface in the microwell or transwell insert, mimicking natural particle deposition at the VF mucosal surface. iLEC-based cultures (Transwell_iLEC_ and VF-OOC_iLEC_) were exposed at 0, 50, 100, and 400 µg/mL for 24 hours. iPSC-based cultures (Transwell_iPSC_ and VF-OOC_iPSC_) were exposed at 0 and 100 µg/mL for 24 hours. All exposure experiments included vehicle controls (0 µg/mL FAD media without PM_10_) and were performed as independent biological triplicates.

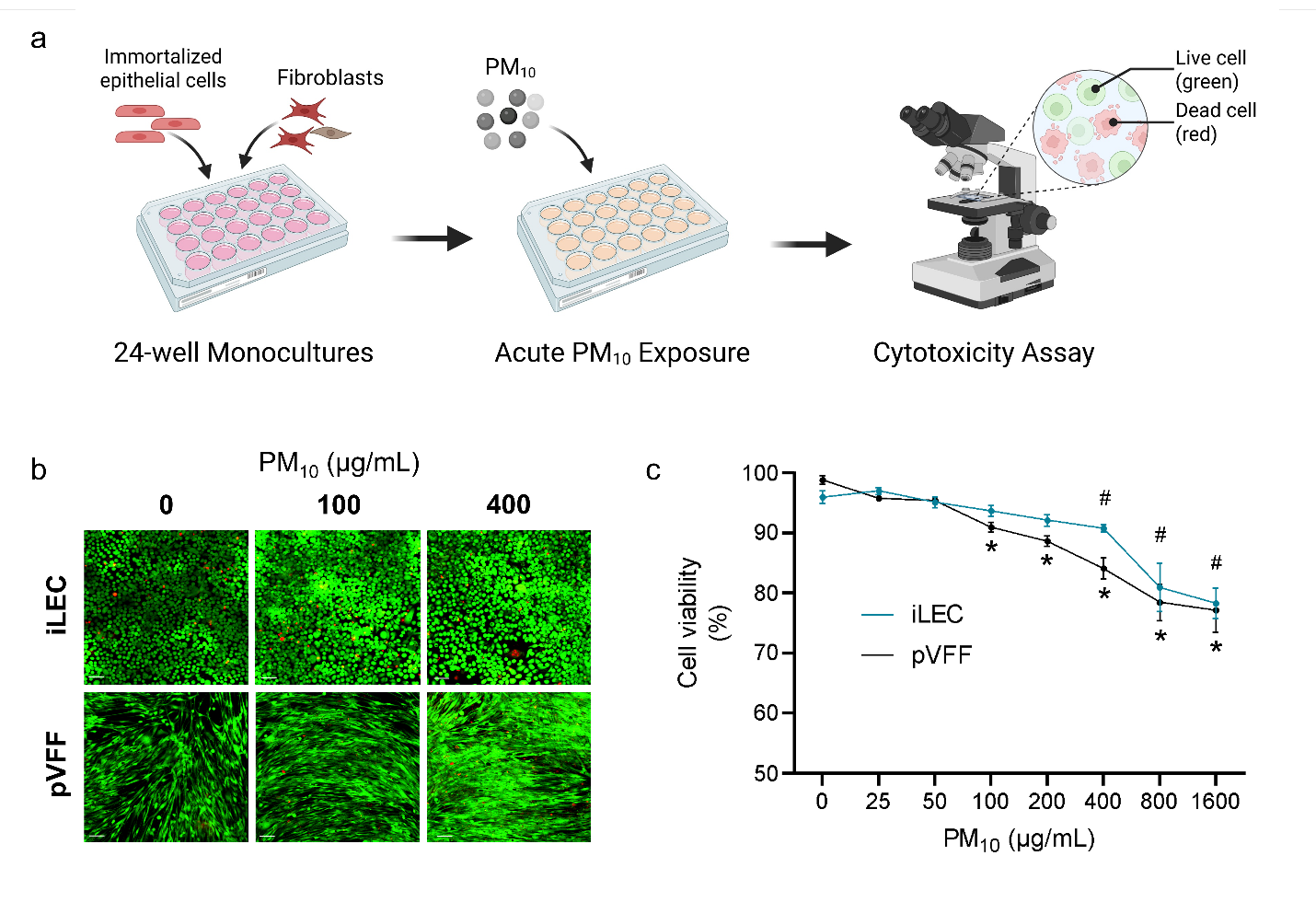
**Figure S2. PM_10_ toxicity screening on pVFF and iLEC monocultures.** (a) Experimental schematic. (b) LIVE/DEAD images at 0, 100, and 400 µg/mL PM_10_. (c) Cell viability versus PM_10_ concentration (0 to 1600 µg/mL). Significant viability reduction versus 0 µg/mL indicated by # (iLEC) and * (pVFF), *p*<0.05. N = 3, mean ± SD.

#### S5. Histology and Immunostaining

**S5.1 Cryosectioning**

Tissue cultures were washed three times in DPBS and fixed in 4% paraformaldehyde (ThermoFisher #J19943K2) for 30 minutes. After washing, microtissue constructs were extracted from culture wells and transferred to cryomolds (Ted Pella #271471) filled with O.C.T. Compound (Fisher #4585). Samples were flash frozen using liquid nitrogen and sectioned vertically at 10 µm thickness with a cryostat (Leica CM1860). Sections were transferred to microscope slides (Fisher #1255015), dried for 1 hour at room temperature, and stored at −80 °C until staining.

**S5.2 H&E Staining**

H&E staining (Abcam #ab245880) was performed following the manufacturer’s protocol. Slides were dipped in hematoxylin, rinsed twice in distilled water, and dipped in bluing reagent. Sections were rinsed in distilled water, briefly immersed in absolute alcohol, and stained with eosin solution. Slides were rinsed again and dehydrated through a graded absolute alcohol series. Processed specimens were mounted using Permount mounting medium (Fisher Scientific #SP15100). Image acquisition was performed using an Axiovert 3 Widefield Microscope (Zeiss) equipped with Zen System software. Images were captured with a 40× objective.

**S5.3 Immunocytochemistry**

Slides were washed in PBS and permeabilized using 0.1% Triton X-100 (Sigma #X100-5ML). After three washes, a blocking buffer comprising 10% goat serum (Abcam #ab7481) and 1% Tween-20 (Abcam #ab128987) in PBS was applied. Samples were incubated with primary antibodies in blocking buffer and then washed with 0.05% Tween-20. If the primary antibody was not fluorescence-conjugated, slides were further incubated in blocking buffer containing fluorescence-conjugated secondary antibody. Samples were counterstained with 1:5000 DAPI (Abcam #ab228549) and mounted with Vectashield Antifade medium (Vector Labs #H-1700). Negative control slides were stained with either secondary antibody only or PBS only. An inverted Zeiss LSM710 confocal fluorescence microscope with a 20× objective was used for image acquisition. Imaris version 7.5.6 software (Bitplane) was used for image analysis. Primary antibodies, dilutions, fluorophores, hosts, and catalogue numbers are listed in Table 1 of the main manuscript.

#### S6. Transmission Electron Microscopy

Samples were washed twice with DPBS and immersion-fixed overnight at 4°C in 2.5% glutaraldehyde and 2% paraformaldehyde in 0.1 M sodium cacodylate buffer (pH 7.4). After washing with sodium cacodylate buffer, samples were post-fixed in 1% osmium tetroxide, dehydrated via a graded ethanol series, washed in propylene oxide, and embedded in Epon 812 epoxy resin under vacuum. Samples were sectioned at 70 nm thickness using a Leica Microsystems EM UC6 Ultramicrotome. Sections were stained with Reynolds lead citrate and 8% uranyl acetate in 50% ethanol to increase image contrast. Images were captured by a Talos F200X STEM transmission electron microscope.

#### S7. Quantitative Real-Time PCR

Microtissue constructs were washed with DPBS and submerged in 100 U/mL of collagenase type I (Gibco #17018029) in HBSS (ThermoFisher #14175095) at 37 °C for 2–3 hours or until gel dissolution. Cells were collected and washed via centrifugation. Supernatant was aspirated, and pellets were snap frozen using liquid nitrogen and stored at −80 °C until RNA extraction.

Total RNA extraction was performed using the RNeasy Micro Kit (Qiagen #74004). Extracted RNA was assessed using a NanoDrop (ThermoScientific, NanoDrop 2000 Spectrophotometer) to determine quantity and purity prior to cDNA synthesis. Samples were further assessed with an Agilent Bioanalyzer to determine RNA integrity prior to qPCR analysis. 500 ng of RNA was reverse transcribed to cDNA using a SuperScript VILO cDNA Synthesis Kit (ThermoFisher #11754050) followed by TaqMan assays (ThermoFisher #4453320).

96-well fast plates (ThermoFisher #4483485) with 1 µL of cDNA per 10 µL reaction mix were run for 40 cycles using a ViiA 7 Real-Time PCR System (ThermoFisher). All cell cultures were performed as independent biological triplicates and run as technical duplicates.

Target genes covered markers of epithelial stratification, tight and gap junctions, mucins, hyaluronan synthases, the mechanosensitive ion channel, and inflammatory cytokines for PM_10_ exposure experiments. Full gene panel and TaqMan assay IDs are listed in Table 1 of the main manuscript. Gene expression was reported relative to GAPDH using the 2^−ΔCt^ method. KRT13 was below the qPCR detection limit in iLEC-based samples and excluded from iLEC analyses. KRT14 was below the detection limit in most iPSC-derived samples and excluded from iPSC analyses.

#### S8. Enzyme-Linked Immunosorbent Assays

Secreted IL-1β (Abcam #ab214025) and TNFα (Abcam #ab181421) proteins were quantified by ELISA. After PM_10_ exposure, supernatant from both the microchannel and microwell was collected and stored at −80 °C until assaying. ELISA plates were prepared according to the manufacturer’s protocol and read using a Tecan Spark Plate Reader (Tecan Life Sciences). ELISA data were normalized to total protein content determined by the Bradford Protein Assay (Bio-Rad Labs #5000002). All samples were cultured as independent biological triplicates and run as technical duplicates.
